# Reduced inhibition of hippocampal adult-born granule cells by parvalbumin interneurons after TBI

**DOI:** 10.64898/2026.09.08.750063

**Authors:** Corwin R. Butler, Connor R. Squellati, Eric Schnell

## Abstract

Traumatic brain injury (TBI) is one of the leading causes of acquired temporal lobe epilepsy. TBI drives hippocampal circuit rearrangements that may contribute to increased seizure risk, such as altered inhibitory circuit function and aberrant post-traumatic neurogenesis. In the hippocampal dentate gyrus, adult-born dentate granule cells (DGCs) acquire inhibitory synaptic inputs from parvalbumin-expressing (PV) interneurons early in their maturation. These inputs are important for circuit integration and feedforward inhibition of these neurons. To test whether DGCs born after TBI have functionally altered PV-mediated innervation, we used genetically modified mice, retroviral vectors, and optogenetics to study adult-born and mature DGCs after TBI. Although DGCs born after TBI acquired inhibitory synaptic inputs during their maturation, PV-mediated inhibition of adult-born DGCs was persistently reduced following TBI. This was not observed in mature granule cells and was not due to TBI-induced changes in PV cell density. This deficit in PV-mediated functional innervation was associated with a transient reduction in release probability at these synapses, which normalized as DGCs matured despite ongoing reduction of functional PV input. Surprisingly, although spontaneous inhibitory postsynaptic currents were reduced for mature granule cells after TBI, these were unchanged in adult-born DGCs. Taken together, these data demonstrate distinct differences in the *de novo* development and maintenance of PV+ synapses in the dentate gyrus after TBI. The addition of neurons with reduced PV+ interneuron-mediated feed-forward inhibition to the dentate gyrus could contribute to hippocampal hyperexcitability after severe brain injury.

## Introduction

Post-traumatic epilepsy (PTE) is a debilitating neurologic condition that can occur after traumatic brain injury (TBI) (Annegers et al., 1998). 25-30% of PTE patients have seizures that are refractory to current medical treatment, and increased brain injury severity is associated with a higher risk of subsequent PTE, which can develop years after the initial injury (Annegers et al., 1998; Temkin, 2009). There is a great need to elucidate the underlying mechanisms associated with epileptogenesis after TBI to support development of improved treatment and prophylactic strategies.

PTE primarily manifests as medial temporal lobe epilepsy (TLE), a form of epilepsy that involves histological and functional changes to the hippocampus (reviewed in Hunt et al., 2013a). One circuit change unique to the dentate gyrus of the hippocampus is aberrant adult neurogenesis, which occurs in both chemoconvulsant models of TLE and animal models of TBI (Butler et al., 2015; Cho et al., 2015; Hendricks et al., 2017; Overstreet-Wadiche et al., 2006; Parent et al., 2006; Villasana et al., 2015). Aberrant neurogenesis after brain injury involves both quantitative changes, through increases in stem cell proliferation and accelerated maturation, and qualitative changes, in patterns of dendritic arborization and neuroblast migration (Carlson et al., 2014; Ibrahim et al., 2016; Kernie et al., 2001; Villasana et al., 2015). TLE is also associated with altered circuit integration of adult-born granule cells, which can affect both the magnitude and timing of their synaptic innervation and the targets of their output projections (Althaus et al., 2019; Hendricks et al., 2017; Overstreet-Wadiche et al., 2006). Ablation of aberrantly generated neurons reduces spontaneous seizure frequency and can improve hippocampal learning and memory in mice with established TLE (Cho et al., 2015), suggesting that these neurons contribute to hippocampal hyperexcitability. Although granule cells born after TBI share morphological similarities to those born after seizure-associated TLE, it is not known whether altered synaptic integration of these neurons might contribute to their post-TBI circuit functions.

Inhibitory interneurons contribute to the sparse activity of the dentate gyrus and prevent excessive hippocampal hyperexcitability (Hunt et al., 2013b; Santhakumar et al., 2000; Sloviter, 1987, 1991; Zhang and Buckmaster, 2009). Dentate interneurons are susceptible to cell death following traumatic brain injury (Butler et al., 2017; Frankowski et al., 2019; Lowenstein et al., 1992), which also coincides with injury-associated changes in the synaptic and tonic inhibition of mature DGCs (Boychuk et al., 2016; Parga Becerra et al., 2021; Zhu et al., 2019). There are several genetically-defined populations of interneurons within the dentate gyrus (reviewed in Houser, 2007) which may be differentially susceptible to injury. For example, somatostatin-expressing interneurons are highly vulnerable to brain injury (Frankowski et al., 2019; Halabisky et al., 2010), while the effects of brain insult on parvalbumin-expressing (PV) interneurons are less consistent (Butler et al., 2022; Folweiler et al., 2020; Frankowski et al., 2019; Zhang and Buckmaster, 2009). To further complicate matters, the effects of brain injury on the density of specific interneuron populations may be only one component of how hippocampal inhibition is altered after TBI, as physiological studies of these cells have uncovered additional abnormalities such as excitation:inhibition ratio changes that could further alter dentate hyperexcitability (Folweiler et al., 2020).

PV interneurons functionally innervate adult-born DGCs early during their maturation, coincident to the timing of aberrant dendritic growth after TBI (Remmers et al., 2020; Vaden et al., 2020; Villasana et al., 2015). As PV cells control feed-forward inhibition in the dentate gyrus [CITE], we investigated whether functional innervation of adult-born DGCs by PV neurons is altered after TBI. We performed the controlled cortical impact (CCI) model of TBI in genetically modified mice, and labeled DGCs born two weeks after TBI with retroviral vectors. We then used whole-cell patch-clamp electrophysiology and optogenetics to assess the functional output from PV+ neurons onto both adult-born DGCs and mature DGCs. We found a selective reduction in the acquisition of PV+ inhibitory inputs by granule cells born after TBI, which was not associated with loss of PV+ inputs by mature DGCs. This selective functional circuit deficit in neurons born during post-traumatic neurogenesis provides an additional mechanism through which these cells might contribute to hippocampal hyperactivity after TBI.

## Materials and Methods

### Animals

All animal care and use procedures were approved by the Institutional Animal Care and Use Committee at the Portland VA. Genetically modified mice selectively expressing Cre in parvalbumin-expressing interneurons (“PV-Cre”) were obtained from the Jackson Laboratory (Bar Harbor, ME; B6.129P2-*Pvalb^tm1(cre)Arbr^*/J; Strain #017320) (Hippenmeyer et al., 2005).

Homozygotic PV-Cre mice were crossed with homozygotic reporter mice expressing Channelrhodopsin2 (“ChR2”) in a Cre-dependent manner (Jackson Laboratory; B6.Cg-Gt(ROSA)*26Sor^tm32(CAG-COP4*H134R/EYFP)Hze^*/J: Stain #024109) (Madisen et al., 2012) to generate double heterozygotic littermates (PV-Cre +/-::ChR2 +/-) used for the experiments in this study. Mice were housed in a 12 hr light/12 hr dark cycle in the AAALAC-approved Portland VA vivarium with food and water provided *ad libitum*. Both male and female mice were used for experiments, with balanced allocation between conditions (for histology experiments: Sham n = 3 male, 4 female and CCI n = 3 male, 4 female; for electrophysiology experiments (5 weeks post-injury): Sham n = 5 male, 3 female and CCI n = 7 male and 4 female; for electrophysiology experiments (8 weeks post-injury): Sham n = 3 male, 5 female and CCI n = 2 male and 5 female; for PV on-cell recordings, Sham n = 1 male, 3 female and CCI n = 2 male, 3 female).

### Controlled cortical impact injury

We used controlled cortical impact (CCI) to induce TBI as previously described (Villasana et al., 2015). Two-month old mouse littermates were randomly allocated to either CCI injury or sham procedure. Mice were anesthetized using isoflurane (2%) mixed with oxygen by spontaneous respiration, and mounted on a stereotaxic apparatus (Kopf Instruments). After sterile preparation and scalp incision, a 5 mm-diameter circular craniotomy was made between lambda and bregma, leaving dura intact, to the right of the midline. Cortical impact was made in the exposed region using a 3-mm-diameter sterile stainless steel tip attached to an electromagnetic impactor (Neuropactor, Neuroscience Tools), set to 4.4 m/sec with an 800 msec dwell time. Deformation depth was proportionally adjusted to account for sex differences in weight; set at 0.9 mm and 0.75 mm in male and female mice respectively. After CCI, the scalp incision was closed by suture and Vetbond glue. Mice recovered in a warm padded chamber and were given soft food soaked with 3.2 mg/ml acetaminophen solution to alleviate pain following surgery. Sham mice received the same treatment (anesthetic, scalp incision/closure, analgesic), with the exception of the craniotomy and impact. Each mouse was individually coded and housed after the procedure. Mice were sacrificed 5 or 8 weeks after sham or CCI procedures.

### Retroviral vector production and labeling of adult-born granule cells

Moloney Murine Leukemia Virus (MMLV)-based retroviral vectors selectively infect mitotic cells, and are an established technique for labeling birthdate-defined adult-born DGCs (van Praag et al., 2002). mCherry-expressing retroviral particles were created using a pRubi-based vector (Addgene #66700) and transfection of GP2 cells as previously described (Luikart et al., 2012), and concentrated to ∼1 x 10^7^ particles/ml using RetroX Concentrator (Takara Bio, Clontech). Retroviral particles were injected in a subset of mice 2 weeks after sham or CCI injury, using coordinates to target both blades of the dentate gyrus in the dorsal hippocampus (from bregma; X = ±1.1, Y = -1.9, and Z = -2.3 and -2.5mm). A volume of 1 µL was injected at each site at 0.25 µL/min (2 injections per hemisphere) using a Hamilton syringe and 30g needle. After each injection, the needle remained in place for at least one minute before being slowly removed. The scalp incision was closed with surgical glue and mice received oral acetaminophen mixed with food for three days as an analgesic. Mice were monitored every 24 hrs for 3 days after injection.

### Electrophysiology

Acute hippocampal slices were prepared from mice 5 or 8 weeks after sham or CCI procedure. Mice were anesthetized using isoflurane by respiration, followed by 2% avertin (0.8 ml, i.p.). Once terminally anesthetized, mice were transcardially perfused using an ice-cold choline chloride-based solution containing the following (in mM): 110 choline Cl, 25 NaHCO_3_, 1.25 NaH_2_PO_4_, 7 MgCl_2_, 0.5 CaCl_2_, 2.4 KCl, 10 D-glucose, and 1.3 Na-ascorbate, with osmolarity ∼320mM. Brains were blocked and glued to a sectioning stage, after which 300 μm coronal hippocampal slices were cut in the same ice-cold choline chloride-based solution on a Leica VT1200S Vibratome. Sections contralateral and ipsilateral to the injury/sham treatment were tracked by isolating them within the holding chamber. Brain slices recovered in artificial cerebrospinal fluid (ACSF; external solution) at 34°C for 30 min, before being transferred to room temperature. ACSF contained the following (in mM): 125 NaCl, 25 NaHCO_3_, 3 KCl, 1.25 NaH_2_PO_4_, 2.0 CaCl_2_, 1.0 MgCl_2_, and 25 D-glucose bubbled with 95% O_2_-5% CO_2_, with osmolarity adjusted to 300-305mM. To isolate inhibitory synaptic signals, 10µM NBQX (Abcam, ab120046) was added to the ACSF during all recordings.

In the recording chamber, slices were continuously perfused with oxygenated ACSF, and individual fluorescent (adult-born) granule cells or ChR2-YFP-expressing PV neurons were identified using combined fluorescence/differential interference contrast imaging, while non-fluorescent mature granule cells were selected from the outer third of the granule cell layer via differential interference contrast imaging. Recording pipettes for mature granule cells and ChR2-expressing PV neurons were pulled from borosilicate glass (TW150F; World Precision Instruments; Claremont, CA) with a P-87 puller (Sutter Instruments, Novato, CA), while recording pipettes for fluorescent (adult-born) DGCs were pulled from leaded borosilicate glass (PG10165; World Precision Instruments; Claremont, CA) with a PC-10 puller (Narishige International; Amityville, NY) to facilitate higher seal resistances. The intracellular solution for voltage clamp recordings contained in (mM): 113 Cesium-Gluconate, 8 NaCl, 10 EGTA, 10 HEPES, 1 MgCl_2_, 1 CaCl_2_, 0.3 MgATP and 2 NaGTP, with final pH balanced to 7.3. Open tip series resistance was 4-7 MΩ for DGC recordings and 3-5 MΩ for on-cell recordings from PV neurons. High-resistance seals were made from visually-identified cells, and whole-cell recordings were obtained by applying brief suction. Cells were voltage-clamped at -70 mV for 5-10 min to allow equilibration of pipette and intracellular solutions prior to collection of evoked responses. To activate PV inputs onto DGCs, 1 msec pulses of LED light (470 nm, 8 mW/cm^2^) were delivered through a 40x 0.8NA water immersion objective (Olympus) directly over the cell being recorded at 0.1 Hz. Paired pulse stimulation was performed with a 100 msec interstimulus interval; paired pulse ratios were calculated from averages of at least 15-20 sweeps per cell by dividing the amplitude of the second evoked response by the amplitude of the first evoked response. After collection of evoked responses, 1 µM TTX was applied to confirm that evoked responses were action potential-dependent; in a subset of cells, we applied 10 µM SR95531 (Abcam) to confirm that all responses were GABAergic. A hyperpolarizing voltage step (−10 mV) was applied before each sweep to monitor series resistance, cell capacitance and input resistance, and cells were not included if intrinsic measures changed >30% during the recording. 3 week old adult-born DGCs were excluded if input resistance was <700 MΩ. We observed no differences in cell input resistances between experimental conditions within groups of DGCs (Table 1). For on-cell recordings of PV-ChR2+ cells, a high resistance seal was formed and maintained without breaking into the cell to preserve membrane integrity. The pipette solution for on-cell recordings contained in (mM): 130 K-Gluconate, 20 KCl, 10 HEPES, 0.1 EGTA, 4 MgATP, and 0.3 NaGTP. Signals were obtained using an Axopatch 1D amplifier, filtered at 5 kHz, and sampled at 10 kHz using custom-written software in IGORPro (WaveMetrics) via a NIDAQ A/D board. Spontaneous IPSCs were detected and quantified using a slope-based algorithm written in IGORPro.

**Table 1:** Summary data.

| Outcome measure | Sham | CCI Contralateral | CCI Ipsilateral | Significance | Multiple comparisons |
| --- | --- | --- | --- | --- | --- |
| PV-oeIPSC amplitudes in 3 week old DGCs | 169 ± 20 pA | 160 ± 23 pA | 86 ± 11 pA | one-Way ANOVA, F(2,22)=5.918, p=0.0088 | Tukey's, Sham vs CCI Contra p=0.94, Sham vs CCI Ipsi p=0.01, CCI Contra vs CCI Ipsi p=0.03 |
| slPSC Frequency in 3 week old DGCs | 0.18 ± 0.07 Hz | 0.15 ± 0.02 Hz | 0.19 ± 0.04 Hz | one-Way ANOVA, F(2,22)=0.1456, p=0.87 | N/A |
| slPSC Amplitude in 3 week old DGCs | 16.6 ± 1.3 pA | 17.0 ± 1.3 pA | 17.2 ± 2.4 pA | one-Way ANOVA, F(2,22)=0.041, p=0.96 | N/A |
| PV oeIPSC amplitudes in mature DGCs (5 weeks post-injury) | 331 ± 66 pA | 256 ± 28 pA | 301 ± 41 pA | one-Way ANOVA, F(2,38)=0.6649, p=0.52 | N/A |
| slPSC Frequency in mature DGCs (5 weeks post- injury) | 0.52 ± 0.05 Hz | 0.51 ± 0.05 Hz | 0.16 ± 0.03 Hz | one-way ANOVA, F(2,37)=20.29, p<0.0001 | Tukey's, Sham vs CCI Contra p=0.99, Sham vs CCI Ipsi p<0.0001, CCI Contra vs CCI Ipsi p<0.0001 |
| slPSC Amplitude in mature DGCs (5 weeks post- injury) | 21.2 ± 1.8 pA | 20.7 ± 2.0 pA | 21.2 ± 2.2 pA | one-Way ANOVA, F(2,37)=0.0224, p=0.98 | N/A |
| % PV-mediated inhibition of 3 week old adult born DGC vs mature DGC, within animals comparison | 58.8 ±4.2% | 56.2 ±7.0% | 29.8 ±3.7% | one-Way ANOVA, F(2,22)=10.98, p=0.0005 | Tukey's, Sham vs CCI Contra p=0.92, Sham vs CCI Ipsi p=0.0007, CCI Contra vs CCI Ipsi p=0.004 |
| PV oelPSC decay tau in 3 week old DGCs | 70.4 ±12.5ms | 88.2 ±16.9ms | 91.7 ±17.2ms | one-Way ANOVA, F(2,22)=0.6085, p=0.55 | N/A |
| PV oelPSC decay tau in mature DGCs (5 weeks post-injury) | 37.4 ±5.5ms | 44.9 ±5.8ms | 37.7 ±4.5ms | one-Way ANOVA, F(2,34)=0.6433, p=0.53 | N/A |
| PV cell density (5 weeks post-injury) | 125 ±16 cells/mm<sup>3</sup> | 140 ±23 cells/mm<sup>3</sup> | 110 ±27 cells/mm<sup>3</sup> | one-way ANOVA, F(2,18)=0.4308 p=0.66 | N/A |
| ChR2-expressing PV neurons (5 weeks post-injury) | 94.8 ±2.0% co-labeling | 91.1 ±2.7% co-labeling | 89.8 ±4.8% co-labeling | one-Way ANOVA, F(2,18)=0.5911, p=0.56 | N/A |
| Optogenetically-evoked APs in ChR2+ PV neurons | 3.2 ±0.5 APs | 3.5 ±0.4 APs | 2.9 ±0.3 APs | one-Way ANOVA, F(2,16)=0.6072, p=0.56 | N/A |
| Paired pulse ratio of PV-to-3 week old DGC inputs | 0.32 ±0.03 PPR | 0.37 ±0.05 PPR | 0.55 ±0.08 PPR | one-Way ANOVA, F(2,20)=4.881, p=0.0188 | Tukey's, Sham vs CCI Contra p=0.78, Sham vs CCI Ipsi p=0.01, CCI Contra vs CCI Ipsi p=0.08 |
| Paired pulse ratio of PV-to-mature DGC inputs (5 weeks post-injury) | 0.40 ± 0.04<br>PPR | 0.38 ± 0.03<br>PPR | 0.39 ± 0.03<br>PPR | one-Way ANOVA,<br>F(2,36)=0.042,<br>p=0.96 | N/A |
| PV oelPSC amplitudes in 6 week old DGCs | 200 ± 20 pA | 169 ± 19 pA | 95 ± 14 pA | one-Way ANOVA,<br>F(2,16)=8.985,<br>p=0.0024 | Tukey's, Sham vs CCI Contra<br>p=0.45, Sham vs CCI Ipsi p=0.002,<br>CCI Contra vs CCI Ipsi p=0.03 |
| sIPSC Frequency in 6 week old DGCs | 0.22 ± 0.04<br>Hz | 0.21 ± 0.03 Hz | 0.24 ± 0.05<br>Hz | one-Way ANOVA,<br>F(2,16)=0.1278,<br>p=0.88 | N/A |
| sIPSC Amplitude in 6 week old DGCs | 20.1 ± 2.5 pA | 19.8 ± 2.8 pA | 20.7 ± 3.1 pA | one-Way ANOVA,<br>F(2,16)=0.025,<br>p=0.97 | N/A |
| PV oelPSC amplitudes in mature DGCs (8 weeks post-injury) | 293 ± 39 pA | 374 ± 59 pA | 324 ± 70 pA | one-Way ANOVA,<br>F(2,29)=0.6149,<br>p=0.55 | N/A |
| sIPSC Frequency in mature DGCs (8 weeks post-injury) | 0.40 ± 0.07<br>Hz | 0.44 ± 0.06 Hz | 0.12 ± 0.03 Hz | one-way ANOVA,<br>F(2,29)=5.632,<br>p=0.0086 | Tukey's, Sham vs CCI Contra<br>p=0.88, Sham vs CCI Ipsi p=0.016,<br>CCI Contra vs CCI Ipsi p=0.015 |
| sIPSC Amplitude in mature DGCs (8 weeks post-injury) | 20.7 ± 1.4 pA | 22.7 ± 2.8 pA | 20.8 ± 3.0 pA | one-Way ANOVA,<br>F(2,29)=0.2315,<br>p=0.79 | N/A |
| % PV-mediated inhibition of 6 week old adult born DGC vs mature DGC, within animals comparison | 75.6 ± 6.5% | 61.5 ± 8.1% | 30.1 ± 5.3% | one-Way ANOVA, F(2,16)=12.16, p=0.0006 | Tukey's, Sham vs CCI Contra p=0.30, Sham vs CCI Ipsi p=0.0005, CCI Contra vs CCI Ipsi p=0.014 |
| Paired pulse ratio of PV-to-6 week old DGC inputs | 0.33 ± 0.05<br>PPR | 0.30 ± 0.05<br>PPR | 0.35 ± 0.08<br>PPR | one-Way ANOVA, F(2,16)=0.1461, p=0.87 | N/A |
| Paired pulse ratio of PV-to-mature DGC inputs (8 weeks post-injury) | 0.38 ± 0.02<br>PPR | 0.29 ± 0.03<br>PPR | 0.33 ± 0.05<br>PPR | one-Way ANOVA, F(2,29)=2.110, p=0.14 | N/A |
| 3 week old DGC cell input resistances | 874 ± 50 MΩ | 860 ± 62 MΩ | 878 ± 65 MΩ | one-way ANOVA F(2,17)=0.1982, p=0.84 | N/A |
| 6 week old DGC cell input resistances | 247 ± 33 MΩ | 261 ± 40 MΩ | 255 ± 36 MΩ | one-way ANOVA F(2,17)=0.1912, p=0.92 | N/A |
| Mature DGC cell input resistances | 254 ± 20 MΩ | 235 ± 31 MΩ | 246 ± 21 MΩ | one-way ANOVA F(2,65)=0.6435, p=0.71 | N/A |

### Perfusion fixation

Perfusion fixation was performed on terminally anesthetized mice after inhaled isoflurane anesthesia followed by i.p. injection of 2% avertin (0.8mL). Mice were then transcardially perfused with 10 mL ice-cold 0.1M phosphate-buffered saline (PBS) followed by 15 mL fixative (4% paraformaldehyde in 0.1M PBS, pH 7.3-7.4). After dissection, brain tissue was post-fixed overnight at 4°C followed by PBS rinsing. Free-floating coronal brain sections were made using a Leica VT1000 vibratome at 100µm thickness.

### Immunohistochemistry

To assess parvalbumin (PV) cell density and reporter expression in the dentate gyrus following CCI injury, free-floating brain sections were permeabilized and blocked using 0.1M PBS with 0.4% Triton-X 100 (PBST) containing 10% goat serum for 1hr at room temperature. Following permeabilization, tissue was stained overnight at 4°C in PBST + 1.5% normal goat serum and the following primary antibodies: anti-parvalbumin (1:1000, guinea pig, Synaptic Systems 195 004), anti-DsRed to amplify the mCherry signal (1:500, rabbit, Tanaka Living Colors 632496), and anti-GFP to amplify the ChR2-YFP signal (1:1000, chicken, Aves Labs, GFP-1020). Sections were then rinsed with PBST 3x 5min and incubated for 4-6hr at room temperature in PBST with 1.5% normal goat serum and secondary antibodies: 1:400 goat anti-chicken 488 (Invitrogen, A11039), 1:400 goat anti-rabbit 568 (Invitrogen, A11011), and 1:400 goat anti-guinea pig 647 (Invitrogen, A11075). After staining, sections were incubated in 1:20K DAPI to stain nuclei, then mounted onto Fisher Superfrost slides with Fluoromount G (Southern Biotech, 0100-01).

### Confocal microscopy and image analysis

Images were acquired using a Yokogawa CSU-W1 spinning disk confocal microscope mounted on a Scientifica Frame with Olympus optics (Intelligent Imaging, Inc). Images were taken with a 20X 0.8NA (air) objective, and PV cell density measurements were made from confocal stacks (∼20µm depth; 0.63µm step) of the dentate gyrus from 3-4 sections per hemisphere per animal taken at 600µm intervals. Cell counts were manually performed in FIJI by a blinded observer using the Cell Counter plugin in FIJI (National Institutes of Health) and divided by the volume of the dentate gyrus granule cell layer in the image to obtain the PV cell density. To determine specificity of reporter expression, the percentage of ChR2-YFP neurons co-labeled in each image was calculated by determining the percentage of ChR2-YFP expressing neurons which also expressed PV. Quantification from individual sections was averaged across the whole mouse for Sham injured mice, and separated into contralateral and ipsilateral hemisphere data for CCI injured mice, to obtain n = 1 per condition per animal.

### Statistics

Graphpad Prism software was used for statistical analysis. All data was first assessed using Shapiro-Wilk normality testing. For normally distributed data, a one-way ANOVA was used to assess differences between sham, contralateral, and ipsilateral conditions with Tukey’s multiple comparisons. When data were not normally distributed, non-parametric testing was used. Significance was set at p<0.05. Data are presented as mean ± SEM in Table 1.

## Results

### Selective reduction of parvalbumin-mediated inhibition after brain injury in adult-born DGCs

To assess the innervation of adult-born DGCs by PV interneurons after traumatic brain injury, we performed controlled cortical impact (CCI) injury or sham surgery on PV-Cre::ChR2 mice. DGCs born two weeks after CCI were labeled using mCherry-expressing retroviral vectors, which selectively infect mitotic cells (van Praag et al., 2002).

When adult-born DGCs were 3 weeks post-mitosis (5 weeks after CCI or Sham injury), we recorded optogenetically-evoked inhibitory postsynaptic responses from PV interneurons (PV oeIPSCs) onto adult-born DGCs using single cell recording techniques. At this stage, these neurons have nearly completed their dendritic outgrowth, but are still refining their synaptic connections (Esposito et al., 2005; Piatti et al., 2011; Toni et al., 2008; van Praag et al., 2002). DGCs born after CCI had reduced PV oeIPSC amplitudes compared to DGCs born after Sham (control) surgery (Figure 1A-D). This effect was only observed ipsilateral to CCI, as adult-born neurons contralateral to the injury had PV oeIPSC amplitudes similar to Sham controls (Figure 1). As expected for evoked IPSCs in immature granule cells (Remmers et al., 2020; Vaden et al., 2020), the decay rate of PV oeIPSCs in 3 week old DGCs was notably slower than the decay of PV inputs onto mature DGCs, but there was no effect of TBI on these kinetics (Table 1). Together, these data suggest a reduction in PV oeIPSCs after TBI that is more likely related to reduced functional innervation of adult-born DGCs by PV neurons, rather than due to changes in the GABA-A receptor composition that have different kinetics.

**Figure 1:**
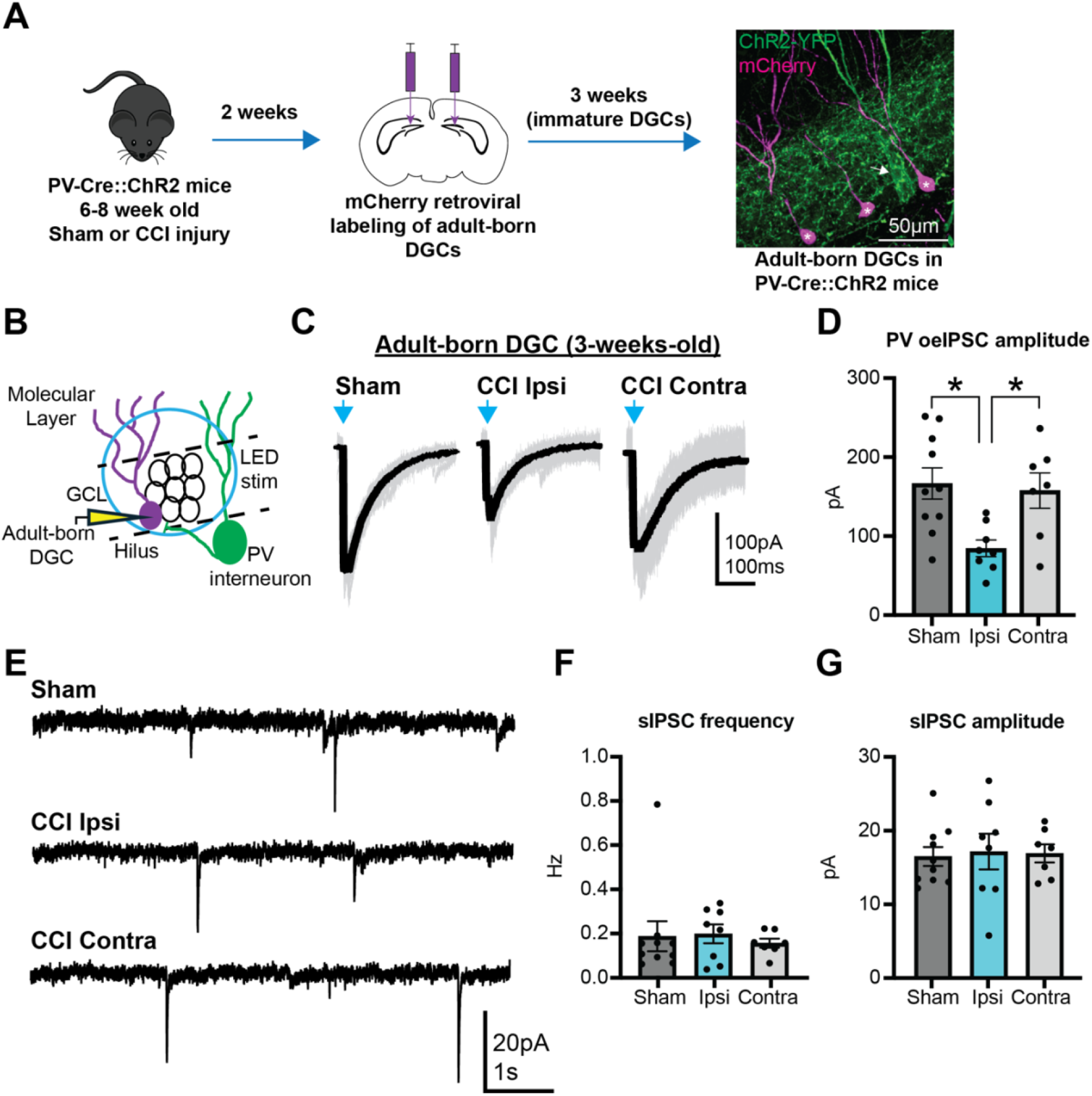
Reduced PV-mediated inhibition of adult-born DGCs after TBI. A) Experimental timeline to test the functional innervation of adult-born dentate granule cells (DGCs) by parvalbumin (PV) interneurons after controlled cortical impact (CCI). In image (right), mCherry-labeled adult-born DGCs (magenta) are noted with asterisks (*); arrow shows a PV neuron expressing Channelrhodopsin2 (ChR2, green) B) Recording configuration schematic. Whole cell recordings from mCherry-labeled adult-born granule cells were obtained while PV interneuron and their axons were optogenetically stimulated with blue light. C) Representative LED light-evoked currents in adult-born DGCs following Sham or CCI injury; blue arrow indicates time of light stimulation. Single traces (grey) shown with overlaid averaged trace (black). D) Optogenetically-evoked PV-mediated inhibitory postsynaptic current (PV oeIPSC) amplitudes recorded from adult-born DGCs in the ipsilateral are reduced after CCI relative to Sham controls (Sham n = 10 cells/ 4 mice; CCI Ipsi, n = 8 cells/ 6 mice; CCI Contra, n = 7 cells/ 6 mice; * = p < 0.05). E) Representative recordings of spontaneous inhibitory postsynaptic currents (sIPSC) in adult-born DGCs following Sham or CCI injury. F) There was no change in sIPSC frequency between groups (n.s. all comparisons). G) There was no change in sIPSC amplitude between groups (n.s. all comparisons).

CCI injury in mouse models causes loss of GABAergic interneurons, and this is paralleled by a reduced spontaneous inhibitory post-synaptic current (sIPSC) frequency in mature DGCs, suggesting an overall reduction in synaptic inhibition due to interneuron loss (Boychuk et al., 2016; Butler et al., 2016; Frankowski et al., 2019; Golub and Reddy, 2022; Hunt et al., 2011; Parga Becerra et al., 2021). However, a similar reduction in sIPSCs following brain injury was not observed in adult-born DGCs identified using genetic labeling strategies (Villasana et al., 2015). To investigate whether these previously-reported differences in overall inhibitory synaptic function between adult-born and mature DGCs after CCI might be technique-related, we measured sIPSC frequencies and amplitudes in 3 week old adult-born DGCs after Sham or CCI injury. We did not observe a difference in mean sIPSC frequency or amplitude between 3 week old adult-born DGCs from each group (Figure 1E-G), suggesting that although PV-mediated inhibition of adult-born DGCs was reduced after TBI, this was not associated with an overall reduction in GABAergic innervation of these immature neurons.

In striking contrast to the findings in adult-born DGCs, there was no difference in PV oeIPSC amplitudes in mature DGCs from CCI or Sham control groups (Figure 2A-C). In these same mature DGCs, however, sIPSC frequency was reduced in DGCs ipsilateral to CCI injury (Figure 2D-F), similar to previous reports (Boychuk et al., 2016; Hunt et al., 2011; Mtchedlishvili et al., 2010; Parga Becerra et al., 2021). Thus, the effects of CCI on PV oeIPSCs and overall sIPSC frequency depended greatly on the maturational stage of the cell. Despite the same post-injury environment, adult-born DGCs demonstrated a selective reduction of PV oeIPSCs without a change in sIPSCs, while mature DGCs had reduced sIPSCs without a change in functional

**Figure 2:**
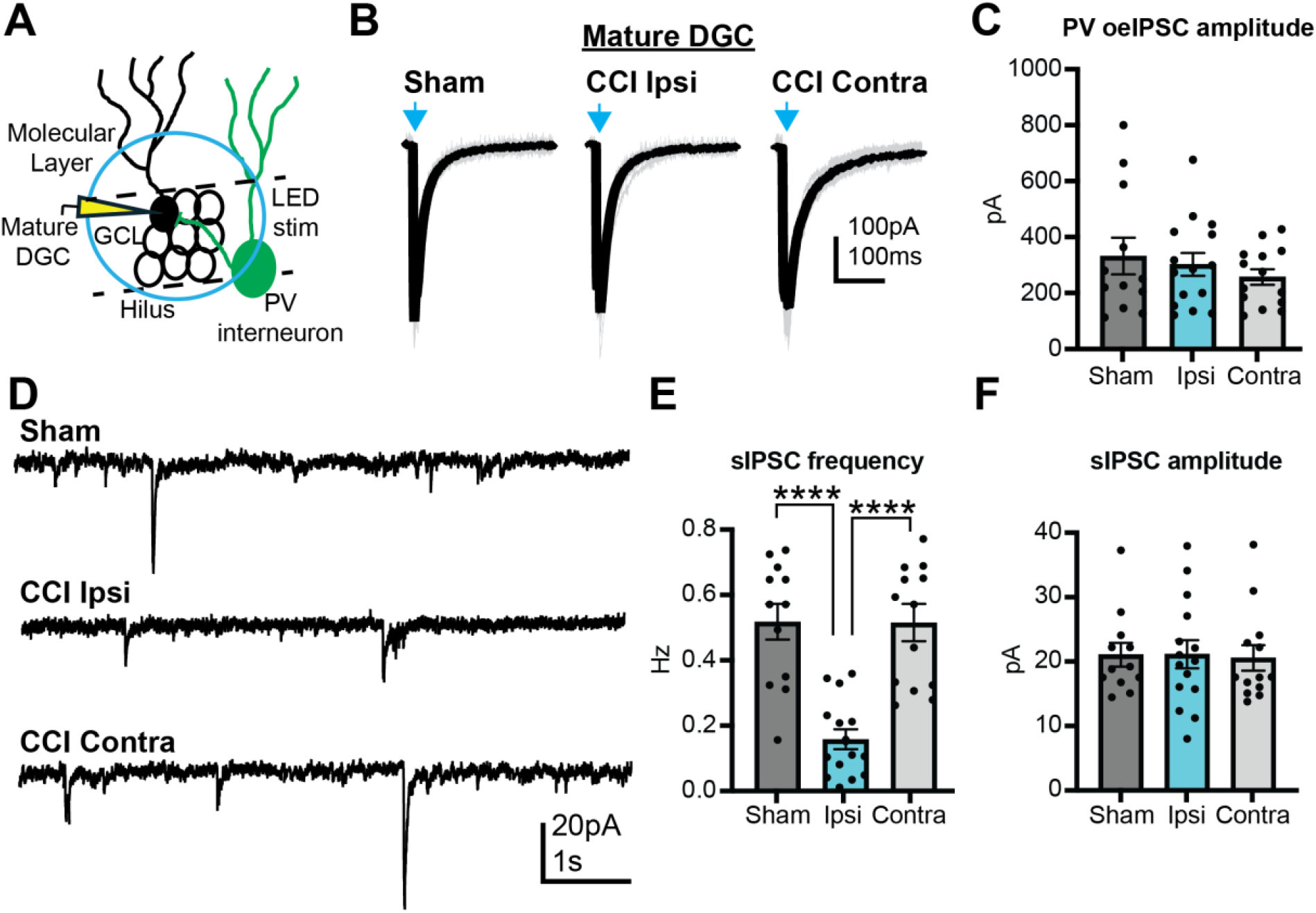
PV-mediated inhibition of mature DGCs is preserved after TBI. A) Recording configuration schematic. LED light-evoked currents were recorded from mature DGCs during optogenetic activation of ChR2-expressing PV interneurons. B) Representative light evoked currents in mature DGCs following Sham or CCI injury; blue arrow indicates time of light stimulation. Single traces (grey) shown with overlaid averaged trace (black). C) Amplitude of optogenetically-evoked PV-mediated inhibitory post-synaptic current (PV oeIPSC) onto mature DGCs is not different between groups (Sham n = 12 cells/ 8 mice; CCI Ipsi n = 15 cells/ 11 mice; CCI Contra, n = 14 cells/ 9 mice; n.s all comparisons). E) Representative recordings of spontaneous inhibitory postsynaptic currents (sIPSCs) in mature DGCs following Sham or CCI injury. F) sIPSC frequency is reduced in mature DGCs ipsilateral to CCI injury compared to Sham and CCI contralateral conditions (****= p < 0.0001). G) No change in sIPSC amplitudes between groups (n.s. all comparisons).

#### PV-mediated innervation

To account for potential variability between groups arising from differences in CCI injury magnitude or slice preparation between animals, we analyzed PV-mediated inhibition in a subset of cells for which each adult-born DGC could be directly compared to a within-animal mature DGC from the same hemisphere. Consistent with the prior analysis (Figure 1), the relative PV-mediated inhibition of adult-born DGCs ipsilateral to CCI injury was substantially smaller than the relative inhibition of adult-born DGCs from sham injured or contralateral to CCI injury (Supplemental Figure 1A-C). In each comparison, PV-mediated inhibition of adult-born DGCs was smaller in amplitude than the PV-mediated inhibition of mature DGCs from the same animal, as has previously been demonstrated for young adult-born DGCs still undergoing synapse acquisition (Groisman et al., 2023); however, this difference was amplified in immature DGCs ipsilateral to CCI. Thus, even when considering potential variability between animals, PV inputs to adult-born DGCs were consistently reduced ipsilateral to CCI injury.

A reduction in inhibitory innervation by PV cells could be caused by injury-induced loss of PV+ interneurons, which occurs in chemoconvulsant epilepsy models and some, but not all, laboratories have observed after TBI (Bouilleret et al., 2000; Folweiler et al., 2020; Frankowski et al., 2019; van Vliet et al., 2004). To determine whether our CCI model caused PV interneuron loss, we measured PV+ cell density in the dentate gyrus of mice following Sham or CCI injury using immunohistochemistry. We did not observe a difference in PV cell density following CCI (Figure 3A,B). Thus, together with the lack of difference in PV oeIPSCS in mature DGCs from Sham and CCI mice, the selective reduction in PV-mediated inhibition of adult-born granule cells ipsilateral to CCI was specific to the innervation of adult-born cells and not explained by a widespread loss of PV inputs.

**Figure 3:**
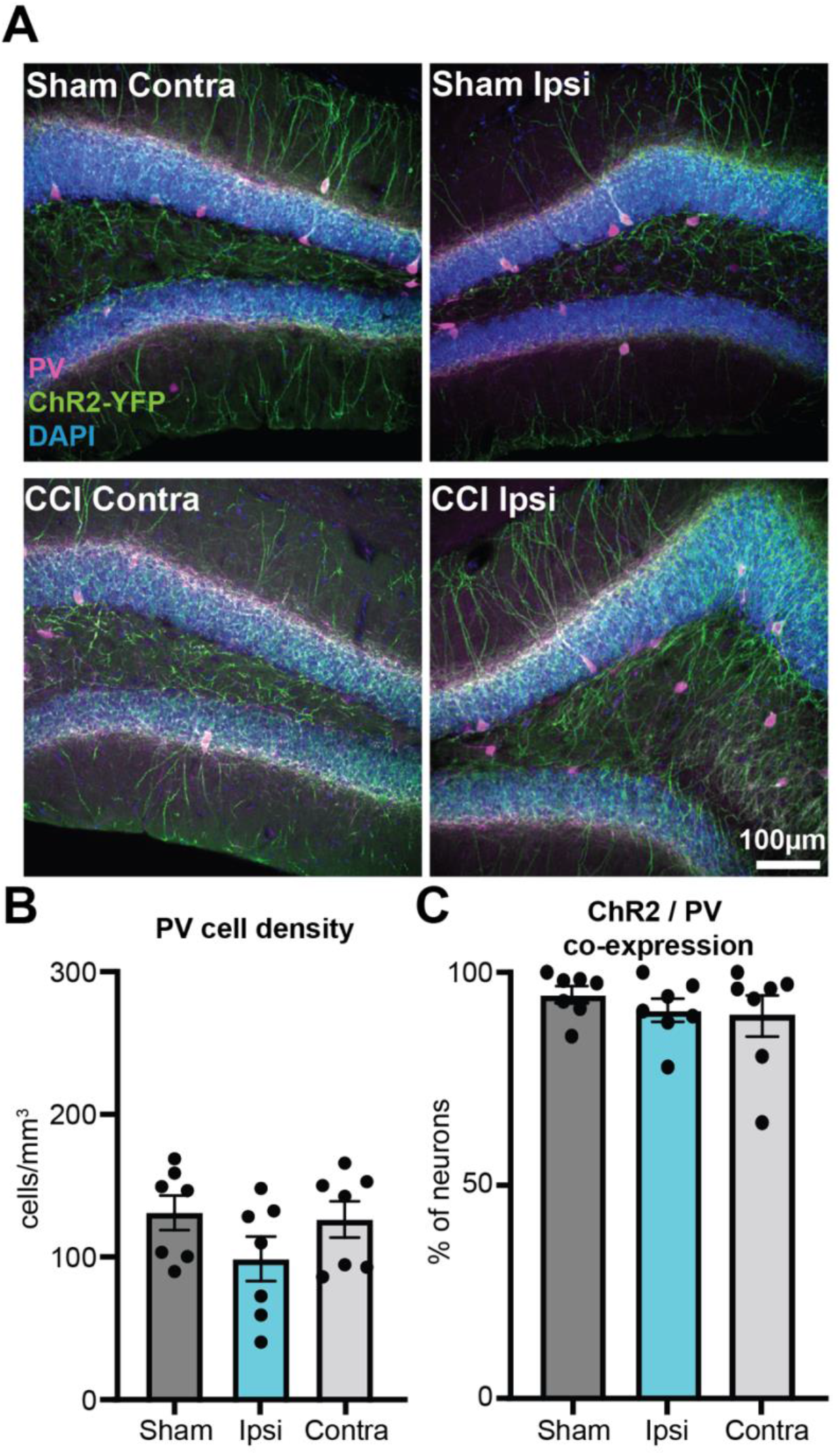
Dentate gyrus PV cell density is unchanged after CCI injury. A) Representative images of stained dentate gyrus sections, showing parvalbumin (PV, magenta), channelrhodopsin2-YFP (ChR2, green), and nuclei (DAPI, blue) in Sham and controlled cortical impact (CCI)-injured mice B) PV cell density in the dentate gyrus is unchanged following CCI injury (Sham, n = 7 mice; CCI hemispheres analyzed separately, n = 7 mice; n.s., all comparisons). C) The percentage of ChR2-positive neurons co-expressing PV is also not different between conditions (n.s., all comparisons).

To further examine potential technical reasons that might explain differences between CCI and sham mice, we examined ChR2 reporter expression and efficacy in PV neurons between conditions. We did not observe a difference in the density of ChR2-expressing PV neurons after CCI injury, indicating that similar numbers of PV cells were expressing ChR2 in each condition (Figure 3C). Additionally, there was no difference in our ability to optogenetically activate PV cells between groups (Supplemental Figure 2). Thus, neither differences in ChR2 expression nor differences in PV cell responsiveness could account for the selective deficit in PV oeIPSCs in adult-born cells after CCI, as similar numbers of PV cells were being optogenetically activated in each condition.

### Presynaptic dysfunction of PV inputs onto adult-born DGCs after TBI

The selective reduction of PV-mediated inputs to adult-born DGCs after brain injury could be due to postsynaptic effects (e.g., fewer GABA receptors) or presynaptic effects (e.g., lower release probability). Presynaptic GABA release from PV-derived synapses onto DGCs is reduced in chemoconvulsant and genetic epilepsy models (Rossignol et al., 2013; Zhang and Buckmaster, 2009), but has not previously been examined after TBI. To examine the presynaptic function of PV cells following CCI injury, we measured paired pulse ratios (PPRs) in both mature and 3-week-old adult-born DGCs from sham and CCI injured mice.

PV oeIPSCs in 3 week old DGCs demonstrated an increase in PPR ipsilateral to CCI, indicating a reduction in presynaptic release probability (Figure 4). This was again selective to adult-born DGCs ipsilateral to CCI, as PV oeIPSCs contralateral to CCI had PPRs similar to those from Sham mice (Figure 4). Consistent with the lack of change in PV oeIPSC amplitudes, there was no difference in PV oeIPSC PPRs recorded from mature DGCs between conditions (Figure 4). These data suggest that the reduction in PV-mediated synaptic inhibition of adult-born DGCs after CCI injury is at least in part mediated by presynaptic dysfunction of these inputs.

**Figure 4:**
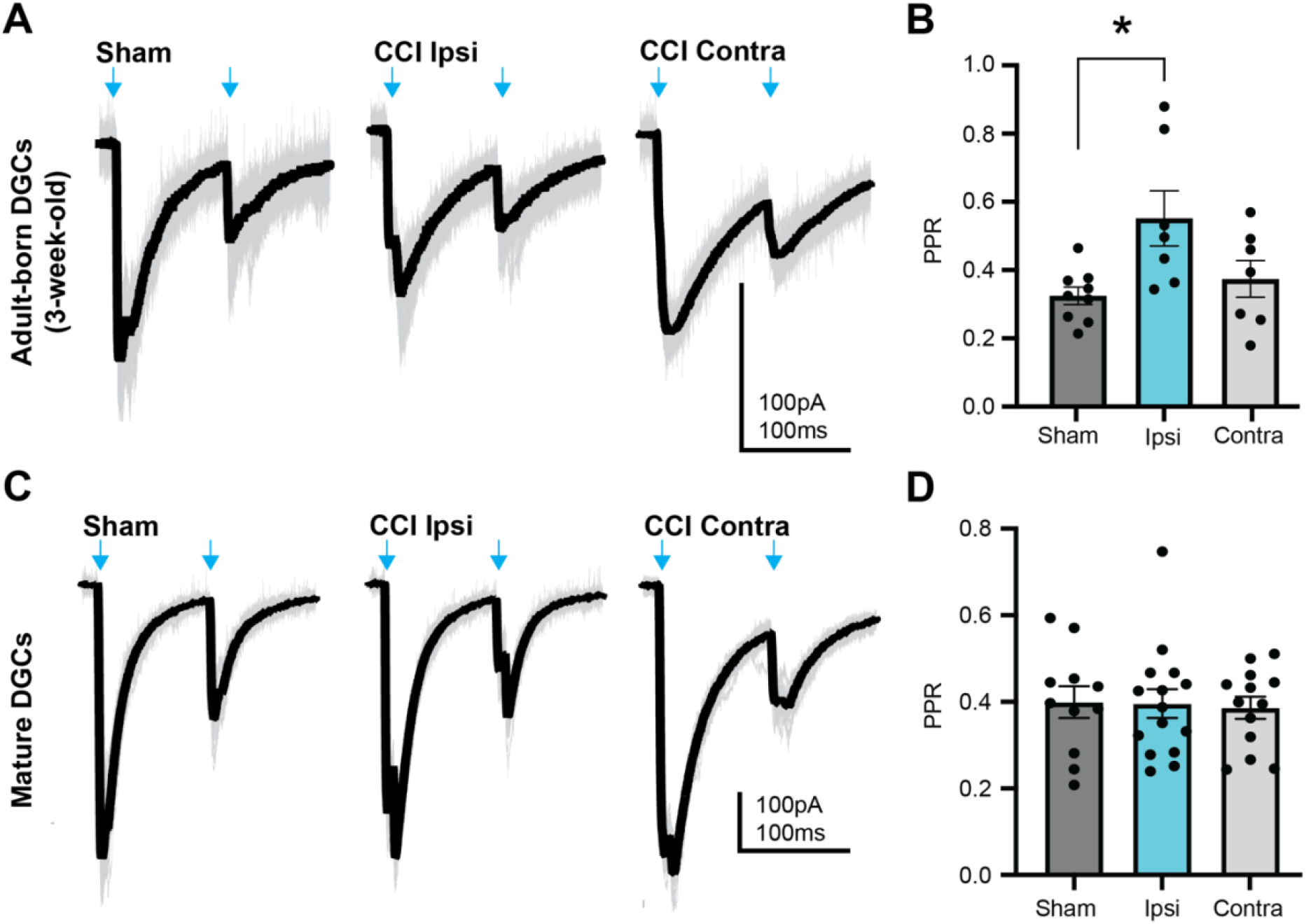
Altered presynaptic function of PV synapses onto 3 week old DGCs born after TBI. A) Representative paired-pulse PV oeIPSCs from adult-born (3 week old) dentate granule cells (DGCs) (Sham n = 10 cells/ 4 mice, CCI Ipsi n = 8 cells/ 6 mice, CCI Contra n = 7 cells/ 6 mice). Blue arrow indicates time of light stimulus. B) Paired pulse ratio of PV-to-adult-born DGC responses is elevated ipsilateral to controlled cortical impact (CCI) injury relative to Sham controls (*, p < 0.05). C) Representative paired-pulse PV oeIPSCs in mature DGCs (Sham n = 12 cells/ 8 mice, CCI Ipsi n = 15 cells/ 11 mice, CCI Contra n = 14 cells/ 9 mice). Blue arrow indicates time of light stimulus. D) Paired pulse ratios of PV-to-mature DGC responses are unchanged following CCI injury (n.s. for all comparisons).

### Persistent reduction in PV-mediated innervation of adult-born DGCs following TBI

Reduced PV-mediated inhibition of immature adult-born DGCs early after brain injury could reflect a transient delay in the presynaptic development of PV-derived inputs, which recovers as granule cells mature. To examine this possibility, we recorded from retrovirus labeled granule cells born 2 weeks after TBI at a later stage (8 weeks after CCI or Sham injury). At this timepoint, these adult-born neurons are 6 weeks post-mitosis and characterized by functionally mature inhibitory synaptic innervation (Laplagne et al., 2006).

Similar to our observations in immature (3 week old) adult-born DGCs, PV-mediated input to 6 week old adult-born DGCs was reduced ipsilateral to CCI injury (Figure 5A-C). Again, sIPSC frequency and amplitude were unchanged in these adult-born DGCs after brain injury, despite the selective reduction in PV-mediated inhibition (Figure 5D-F). This indicates that the deficiency in PV-mediated functional innervation of immature adult-born cells is sustained as these cells mature into the hippocampal circuit.

**Figure 5:**
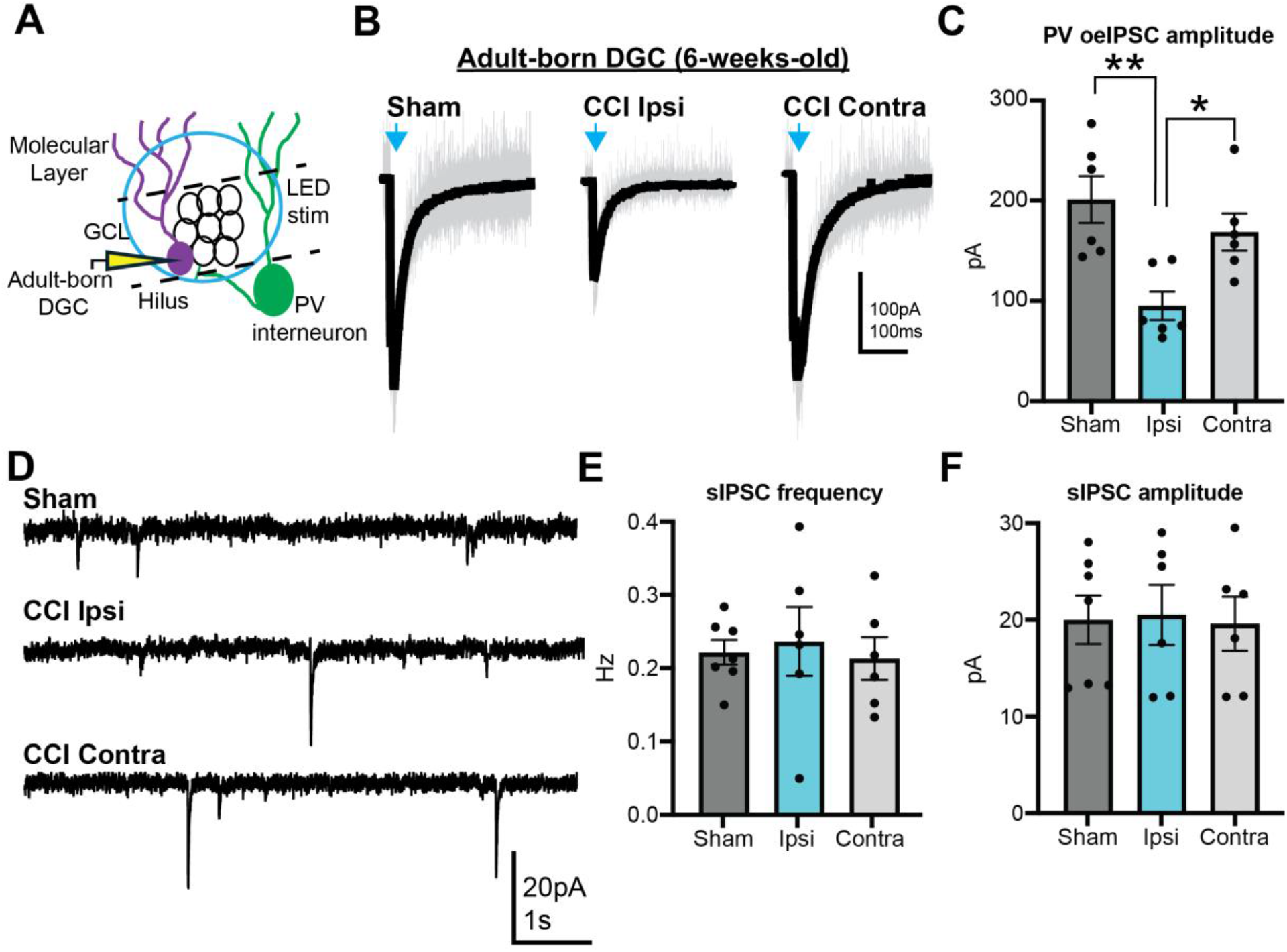
Persistent reduction of PV-mediated inhibition of adult-born DGCs born after TBI. A) Recording schematic. B) Representative PV oeIPSCs recorded from 6 week old adult-born DGCs following Sham or CCI injury; blue arrow indicates time of LED light stimulation. C) PV oeIPSC amplitude in adult-born DGCs (6 weeks old) is reduced in the ipsilateral hemisphere after controlled cortical impact (CCI) injury relative to Sham controls (Sham n = 7 cells/ 6 mice, CCI Ipsi n = 6 cells/ 4 mice, CCI Contra n = 6 cells/ 5 mice; * = p < 0.05). D) Representative recordings of spontaneous inhibitory postsynaptic currents (sIPSC) in 6 week old adult-born DGCs following Sham or CCI injury. E) There was no change in sIPSC frequency between groups (n.s. all comparisons). F) There was no change in sIPSC amplitude between groups (n.s. all comparisons).

As in prior experiments performed at the earlier timepoint, we performed the same experiments in nearby mature granule cells in the outer third of the granule cell layer to further characterize PV circuit function following TBI. Similar to our prior observations 5 weeks after injury, we did not see a difference in PV-mediated inputs to mature DGCs 8 weeks after CCI injury (Figure 6A-C). Despite the lack of change in PV oeIPSCs, sIPSC frequency was again reduced in mature DGCs ipsilateral to CCI, with no change in sIPSC amplitude at this later timepoint (Figure 6 D-F). This indicates that although overall inhibition of mature DGCs is reduced after CCI injury, PV-mediated innervation appears to be preferentially maintained.

**Figure 6:**
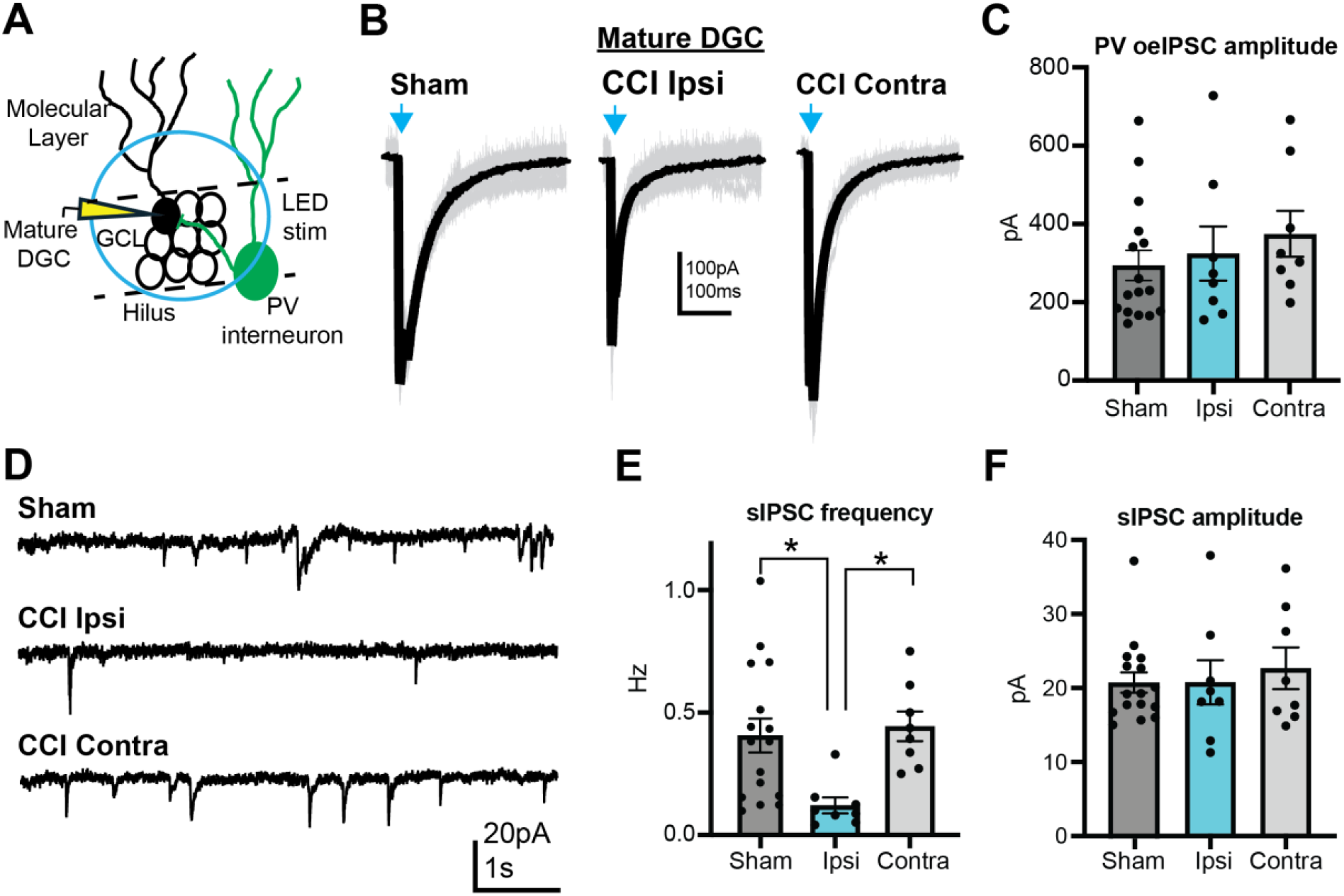
PV-mediated innervation of mature DGCs 8 weeks after TBI is unchanged despite reduced sIPSC frequency. A) Recording schematic. B) Representative PV oeIPSCs in mature DGCs following Sham or CCI injury; blue arrow indicates time of light stimulation. C) PV oeIPSC amplitudes in mature DGCs are not different between groups (Sham n = 16 cells/ 8 mice, CCI Ipsi n = 8 cells/ 7 mice, CCI Contra n = 8 cells/ 7 mice; n.s all comparisons). D) Representative sIPSC traces recorded from mature DGCs from sham and post-CCI tissue. E,F) Mean sIPSC frequency is reduced in mature DGCs ipsilateral to CCI with no change in sIPSC amplitude (* p < 0.05).

When controlling for potential between-animal differences in CCI injury magnitude or slice preparation, at 8 weeks after injury there remained a significant relative reduction in PV-mediated inhibition of adult-born DGCs ipsilateral to CCI injury (Supplemental Figure 1D,E). When combined with our observations at 5 weeks after CCI, this demonstrates that the reduction in PV-derived functional innervation of adult-born DGCs after brain injury is selective and sustained, while PV-derived innervation of mature DGCs continues to be maintained at later stages after brain injury.

To examine presynaptic function of PV inputs after CCI at this later stage, we again used paired-pulse stimulation to examine GABA release from PV neurons onto mature and adult-born DGCs. Interestingly, paired pulse ratios of PV inputs onto 6 week old DGCs were not different between experimental groups (Figure 7A-C). Similar to our prior observations, we did not see a change in the paired pulse ratio of evoked PV responses onto mature DGCs (Figure 7D,E). This would suggest that as DGCs born after CCI mature and more fully integrate into the hippocampal circuit, the presynaptic function of PV neurons onto these newly generated cells normalizes. Despite this normalized presynaptic function, however, PV-mediated inhibition of adult-born DGCs remain reduced, suggesting that mechanisms other than reduced GABA release now account for this reduced functional innervation.

**Figure 7:**
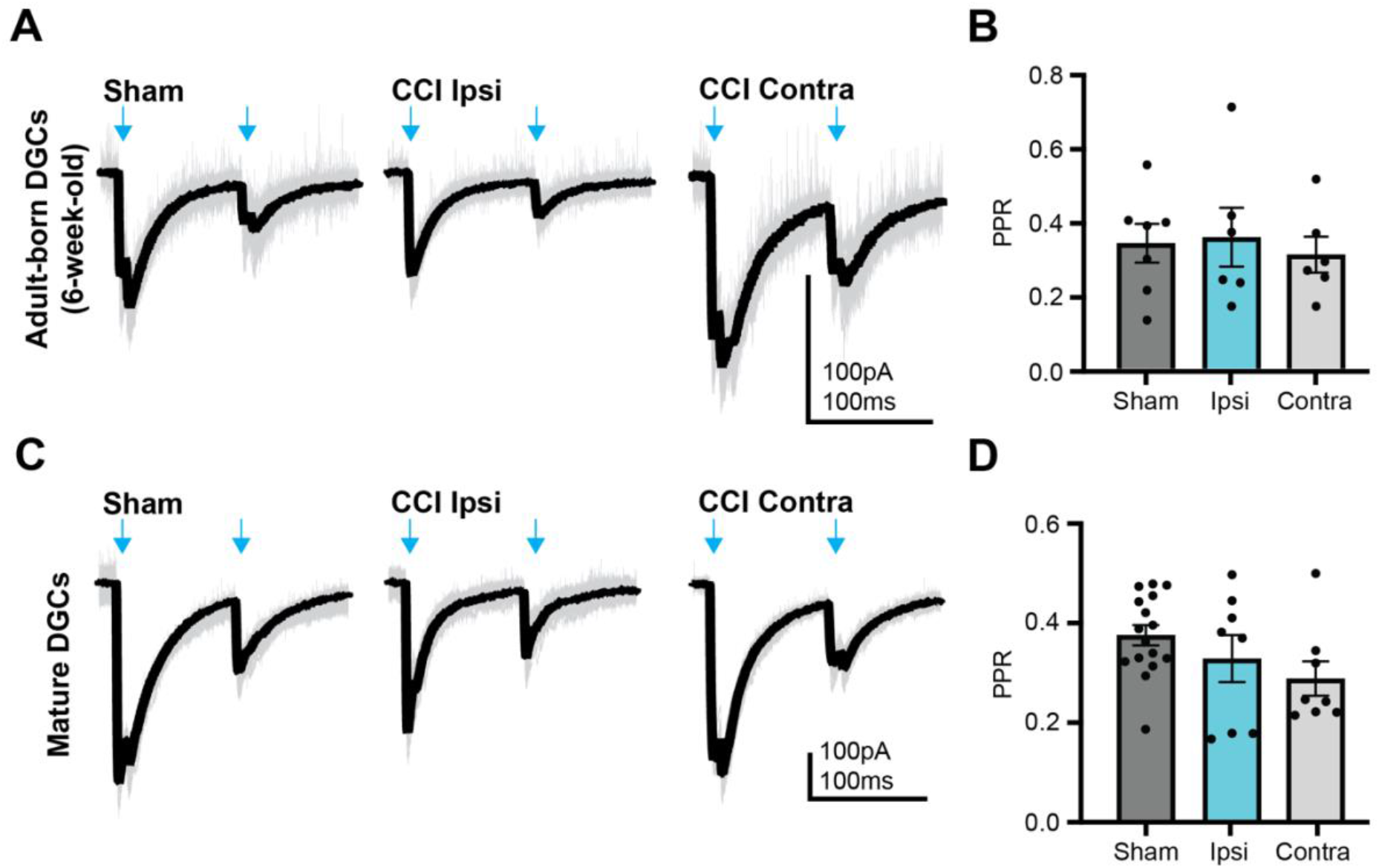
Pre-synaptic release probability of PV oeIPSCs normalizes as adult-born DGCs mature following TBI. A) Representative PV oeIPSCs recorded from 6 week old adult-born dentate granule cells (DGCs) during paired pulse light stimulation; blue arrow indicates time of light stimulus. B) Paired pulse ratio (PPR) of PV-to-adult-born DGC responses is not different between groups (Sham n = 7/ 6 mice cells, CCI Ipsi n = 6 cells/ 4 mice, CCI Contra n = 6 cells/ 5 mice; n.s. for all comparisons). C) Representative PV oeIPSCs recorded from mature DGCs; blue arrow indicates time of light stimulus. D) PPRs of PV-to-mature DGC responses are not different between conditions (Sham n = 16 cells/ 8 mice, CCI Ipsi n = 8 cells/ 7 mice, CCI Contra n = 8 cells/ 7 mice; n.s. for all comparisons).

## Discussion

As the onset of PTE is often significantly delayed relative to injury, it is critical to understand how hippocampal circuit structure and function change over time, and how these various changes might contribute to increased hippocampal excitability. In preclinical models of TBI, hilar neuron loss and aberrant adult neurogenesis are concurrent with elevated seizure susceptibility, but functional studies of the individual circuit elements are ultimately necessary to define how they are affected by injury. Here, we identify a sustained TBI-associated deficit in inhibitory innervation of adult-born DGCs by a subset of genetically-defined of interneurons, which could contribute to hippocampal hyperexcitability during circuit activity, and the occurrence of seizures in PTE.

### Parvalbumin interneurons and inhibitory circuit dysfunction after TBI

There are many different interneuron subtypes in the hippocampal dentate gyrus, which can be categorized based on protein expression, interneuron structure, or physiological parameters (reviewed in Houser, 2007). The vulnerability of each of these different interneuron populations to TBI could depend on the severity of initial injury, in addition to likely intrinsic differences in susceptibility to trauma between neuron types. As the reduction of PV inputs onto adult-born DGCs was not associated with an overall reduction in sIPSC frequency in these cells, other changes in the GABAergic circuit, such as increased acquisition of synapses from other interneuron subtypes, or enhanced activity in surviving interneuron populations following CCI, could partially compensate for their loss of functional PV-mediated innervation (Boychuk et al., 2022; Hunt et al., 2011; Kang et al., 2022). Alternatively, it is possible that non-PV interneurons are the dominant contributors to sIPSC frequency measured in adult-born DGCs, in which case the isolated loss of PV-mediated synapses would not drive a detectable change in this measure.

Despite the lack of change in overall sIPSC frequency in adult-born DGCs after CCI, however, the loss of PV-mediated inhibition might selectively reduce inhibitory control of these DGCs during higher levels of incoming activity, during which PV-mediated feed-forward inhibition could be required to prevent sustained DGC activation (Espinoza et al., 2018). Additionally, if these adult-born neurons also contributed to recurrent mossy fiber sprouting after CCI (Hendricks et al., 2017; Hunt et al., 2010; Parent et al., 2006), the loss of PV-mediated inhibition of even a subset of DGCs could have substantially increase hippocampal hyperexcitability.

Interestingly while the effects of CCI injury on PV-mediated evoked inhibition and sIPSC frequency in adult-born DGCs were consistent across timepoints, this was not true for the observation of pre-synatic dysfunction which recovered as adult-born DGCs matured in the injured environment. This observation mirrors prior work on synapse development within the hippocampus, including GABAergic synapses in adult-born DGCs (Hsia et al., 1998; Mozhayeva et al., 2002). Although we observed a “normalization” of PPR in adult-born DGCs after CCI when these neurons were more mature, there may be other factors like number of release sites or transmission success which could contribute to this observation (Kang et al., 2022; Zhang and Buckmaster, 2009).

Conversely, functional PV-derived inputs were maintained on mature cells after CCI despite a substantial loss in sIPSCs in those cells, suggesting that loss of inhibition from other interneuron populations, such as somatostatin-expressing interneurons, could play an additional role in the inhibitory circuit remodeling of the dentate gyrus after TBI (Butler et al., 2017; Hunt et al., 2011). While some of the reduction in the inhibition of mature DGCs may be directly related to the loss of specific cell types after TBI (Zhu et al., 2019), in the case of PV-derived inhibition, the deficit in PV innervation of adult-born cells relates selectively to *de novo* synaptogenesis, indicating that cell counts alone do not capture the effects of TBI on dentate function. Together, these findings underscore the importance of circuit-specific functional analyses after TBI to facilitate a full understanding of how brain injury alters hippocampal function.

Why might the PV-mediated functional innervation of adult-born DGCs differ so dramatically from that observed in mature neurons after brain injury? One explanation could result from the distinct mechanisms controlling synapse formation versus synapse maintenance (Bouwman et al., 2004; Lin and Koleske, 2010). Specifically, mechanisms that control the formation of synapses from PV neurons might be negatively impacted in the post-TBI dentate gyrus in a manner that does not extend to loss of previously-existing PV synapses. For example, innervation of adult-born DGCs by PV interneurons is in part driven by the synaptic adhesion molecule, neuroligin-2, and knockdown of neuroligin-2 in adult-born DGCs reduces PV+ synaptogenesis and release probability from PV inputs onto adult-born DGCs (Groisman et al., 2023). This impairment was not observed in distal inhibitory inputs, suggesting that the role of neuroligin-2 is selective for the perisomatic inputs from PV interneurons (Groisman et al., 2023). Currently, it is not known whether TBI selectively alters the expression of inhibitory synaptogenic molecules after TBI in mice, but at least one prior report has demonstrated that the phosphorylation state of neuroligin-2 could be altered after TBI (Lizhnyak and Ottens, 2015), suggesting that both proteomic and transcriptomic analyses of adult-born granule cells could contribute to a molecular understanding of how TBI alters synaptic integration of adult-born neurons. As our understanding of the molecular pathways controlling synaptogenesis and synapse maintenance between specific interneuron types and granule cells increases (Favuzzi et al., 2019; Sanes and Zipursky, 2020), which could include both adhesion molecules and secreted factors (Dickins and Salinas, 2013), it may be possible to one day use this information to restore the appropriate innervation of adult-born DGCs after brain injury.

In the days following TBI in mice, altered levels of neuronal activity can extend well beyond the directly injured area of cortex, and suppression of enhanced activity can reduce aberrant neurogenesis (Tierno et al., 2026; Villasana et al., 2019). As GABAergic synapse development in the brain is regulated by activity (reviewed in Oh and Smith, 2019), altered GABAergic synaptogenesis after brain injury could be driven by changes in activity. Following chemoconvulsant-induced status epilepticus, PV neuron activity is reduced, and increasing activity of these neurons using chemogenetics can reduce seizures (Lee et al., 2024). Notably, reducing global excitability following CCI injury with the GABA-A receptor agonist diazepam reduces aberrant neurogenesis (Villasana et al., 2019), which suggests another mechanism driving aberrant PV-mediated innervation adult-born DGCs after brain injury. While it remains unclear what mechanisms drive differences in PV-to-adult-born DGC synapse generation versus the maintenance of PV-to-mature DGC synapses, continued efforts in this research will not only help us better understand how DGC integration is orchestrated, but may also provide molecular targets for intervention following brain injury to prevent abnormal circuit changes.

### Role of ongoing post-traumatic aberrant neurogenesis in hippocampal circuit dysfunction

Brain injury causes aberrant adult neurogenesis, which alters the number and structure of adult-born neurons (Butler et al., 2015; Niv et al., 2012; Parent et al., 2006; Villasana et al., 2015). The density of immature DGCs changes dramatically during the first weeks following TBI, suggesting dynamic quantitative changes in granule cell neurogenesis. For example, although most young or newly born DGCs generated during the first week after brain injury do not survive (Carlson et al., 2014; Gao et al., 2008), neurogenesis rates return and increase above baseline during the second week after injury (Butler et al., 2015; Dash et al., 2001; Villasana et al., 2015). At remote timepoints, neurogenesis is reduced to much lower levels (Golub and Reddy, 2022), which may relate to stem cell depletion secondary to a large increase in post-injury neurogenesis (Neuberger et al., 2017; Ngwenya and Danzer, 2018).

Although enhanced post-injury adult neurogenesis is associated with improved behavioral recovery in the first few weeks after TBI (Carlson et al., 2014; Kleindienst et al., 2005; Sun et al., 2007; Wu et al., 2018), more recent work has suggests that aberrant integration of adult-born neurons contributes to circuit rearrangements associated with hyperexcitability (Hendricks et al., 2017; Lybrand et al., 2021; Sparks et al., 2020; Villasana et al., 2015). Given that adult neurogenesis is an ongoing process, it is possible that dysfunction associated with aberrant post-injury neurogenesis could compound over time. Although our study only focused on a very specific timepoint after injury, if DGCs born after TBI continue to receive less PV-mediated inhibition, these neurons could accumulate over time in a manner that promotes population hyperexcitability and seizures. Although this could be mitigated by lower rates of neurogenesis at later timepoints (Golub and Reddy, 2022), the persistence of inhibitory dysfunction would still be expected to produce sustained circuit disinhibition.

Another factor that could influence how adult-born DGCs contribute to later circuit dysfunction is the birth date of these neurons relative to the insult. Adult-born DGCs that are immature at the time of injury develop abnormal feed-forward inhibitory microcircuits within the dentate gyrus (Kang et al., 2022), and prior work in chemoconvulsant models has also suggested that DGCs born shortly before status epilepticus might produce enhanced contributions to axonal remodeling (Hendricks et al., 2017; Kron et al., 2010). Future work investigating the functional integration of populations of DGCs born at various timepoints relative to the time of injury will help elucidate how these adult-born DGCs contribute to circuit dysfunction in injured environments.

As changes in individual inputs may be grossly balanced by compensatory changes in other circuits, a complete understanding of how neurons born after injury function in hippocampal circuits will require dissection of these individual circuit elements. The work in this study demonstrates that DGCs born after brain injury manifest a reduction in functional PV-mediated inhibition despite the fact that mature granule cells in the same circuits do not experience the same loss. Since this reduction persists even as these neurons mature into the hippocampal circuit, this may lay the groundwork for latent circuit dysfunction associated with diseases like PTE.

## Acknowledgements

We thank members of the Schnell and Westbrook labs and Dr. Jeffery Boychuk for feedback and support. This work was funded by NIH Grants R01NS126247 (ES), VA I01-BX004938 (ES), VA I01-BX006921 (ES), and VA IK2-BX005761 (CRB). The contents of this manuscript do not represent the views of the US Department of Veterans Affairs or the US government.

---

**Supplemental Figure 1:**
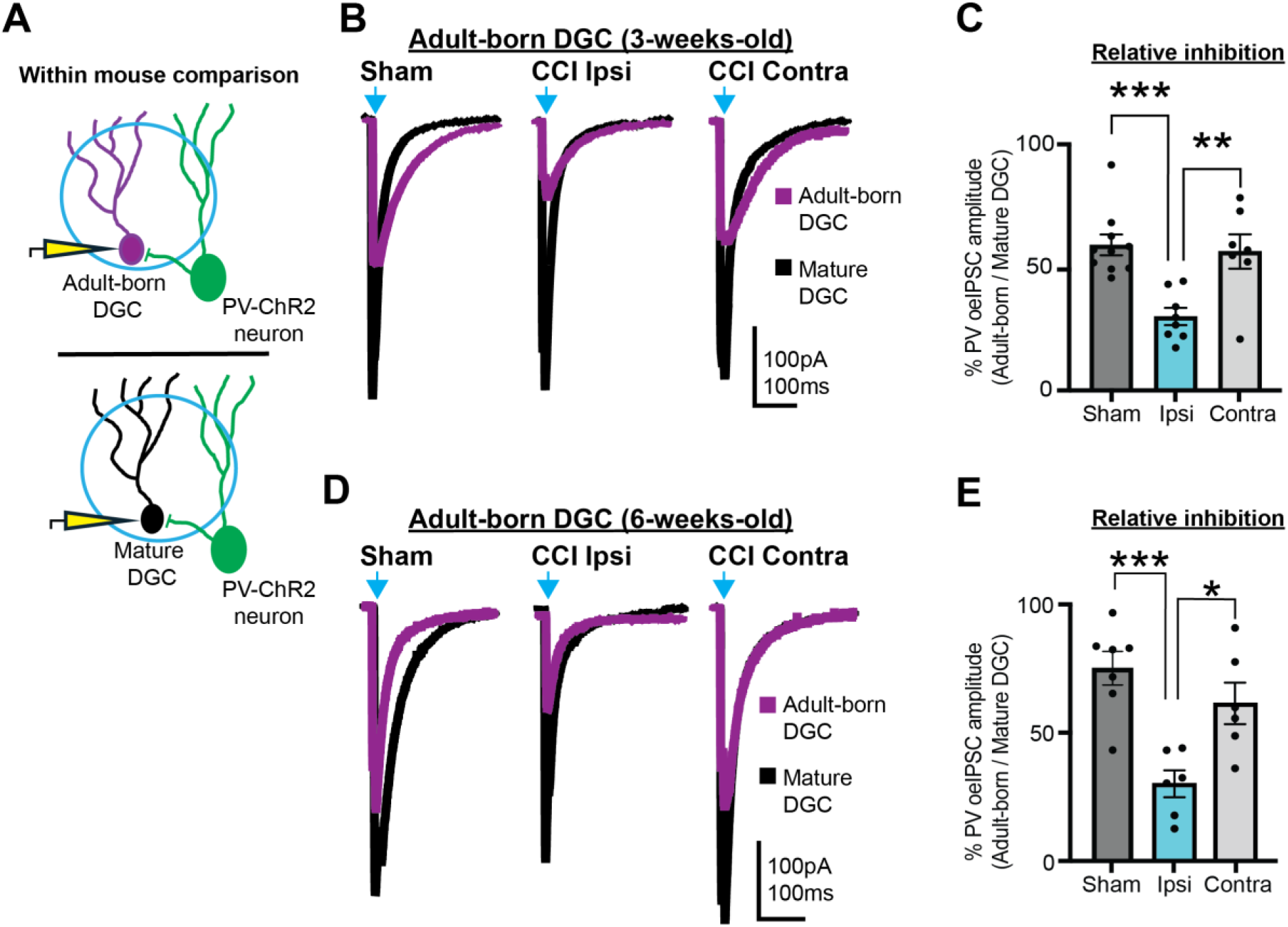
Within-mouse comparisons demonstrate selectively reduced functional innervation of adult-born DGCs by PV cells after TBI. A) Schematic of within-animal comparisons of PV oeIPSCs in adult-born vs mature DGCs from the same animal/hemisphere. B) Representative PV oeIPSCs recorded from 3 week old adult-born (magenta) and mature (black) DGCs following Sham or CCI injury; blue arrow indicates time of light stimulation. Each pair of overlaid traces were recorded from the same animal/hemisphere. C) PV oeIPSC amplitudes in adult-born DGCs were consistently smaller than those recorded from their mature counterparts, but this ratio was further reduced in adult-born cells ipsilateral to CCI relative to Sham and CCI contralateral controls (Sham, n = 10 adult-born/ mature pairs, 6 mice; CCI Ipsi, n = 8 adult-born/ mature pairs, 4 mice; CCI Contra, n = 7 adult-born/ mature pairs, 4 mice; ** p < 0.01, *** p < 0.001). D) Representative PV oeIPSCs recorded from 6 week old adult-born and mature DGCs following Sham or CCI injury; blue arrow indicates time of light stimulation. Each pair of overlaid traces were recorded from the same animal/hemisphere. E) Similar to PV oeIPSCs in 3 week old adult-born cells, relative PV-mediated inhibition of 6 week old DGCs is reduced ipsilateral to CCI relative to Sham and CCI contralateral controls (Sham n = 7 adult-born/ mature cells, 6 mice; CCI Ipsi n = 6 adult-born/ mature cells, 4 mice; CCI Contra n = 6 adult-born/ mature cells, 5 mice; ** p < 0.01, *** p < 0.001).

**Supplemental Figure 2:**
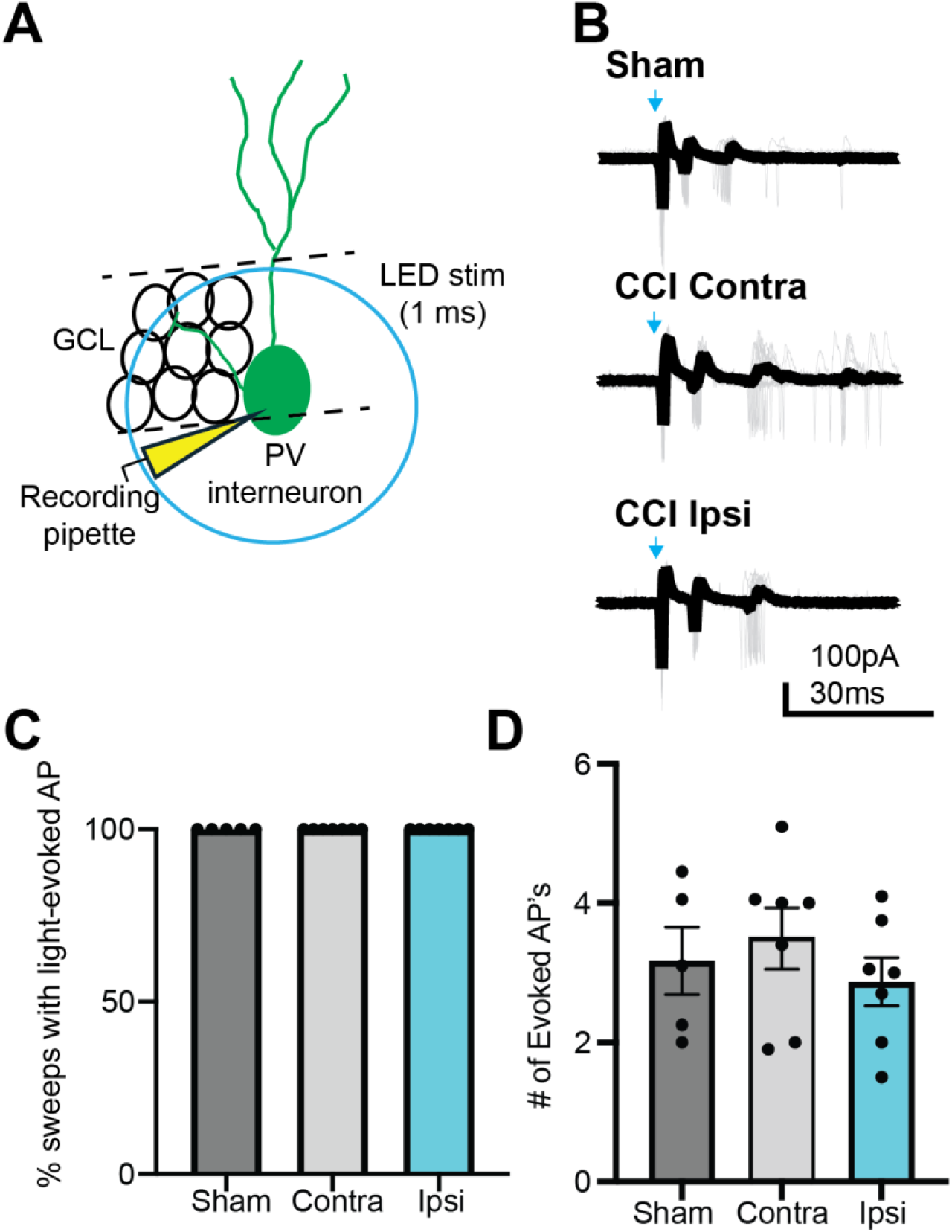
Optogenetic activation of PV neurons is not altered by CCI. A) Recording schematic. Cell-attached recordings were obtained from ChR2-expressing parvalbumin (PV) neurons in the dentate gyrus during 1ms LED light (470nm) stimulation to evoke action potentials (APs). B) Representative optogenetically-evoked APs in PV neurons from Sham and CCI animals following LED stimulation; single sweeps shown in grey with averaged trace superimposed in black. C) ChR2-expressing PV neurons in the dentate gyrus fired at least one AP in response to light stimulation in 100% of sweeps (20 sweeps per neuron) regardless of injury condition. D) The average number of light-evoked APs in PV-ChR2+ neurons was not different between experimental groups (n.s. for all groups).

